# Potential autoimmunity resulting from molecular mimicry between SARS-CoV-2 Spike and human proteins

**DOI:** 10.1101/2021.08.10.455737

**Authors:** Janelle Nunez-Castilla, Vitalii Stebliankin, Prabin Baral, Christian A Balbin, Masrur Sobhan, Trevor Cickovski, Ananda Mohan Mondal, Giri Narasimhan, Prem Chapagain, Kalai Mathee, Jessica Siltberg-Liberles

## Abstract

SARS-CoV-2 causes COVID-19, a disease curiously resulting in varied symptoms and outcomes, ranging from asymptomatic to fatal. Autoimmunity due to cross-reacting antibodies resulting from molecular mimicry between viral antigens and host proteins may provide an explanation. We computationally investigated molecular mimicry between SARS-CoV-2 Spike and known epitopes. We discovered molecular mimicry hotspots in Spike and highlight two examples with tentative autoimmune potential and implications for understanding COVID-19 complications. We show that a TQLPP motif in Spike and thrombopoietin shares similar antibody binding properties. Antibodies cross-reacting with thrombopoietin may induce thrombocytopenia, a condition observed in COVID-19 patients. Another motif, ELDKY, is shared in multiple human proteins such as PRKG1 and tropomyosin. Antibodies cross-reacting with PRKG1 and tropomyosin may cause known COVID-19 complications such as blood-clotting disorders and cardiac disease, respectively. Our findings illuminate COVID-19 pathogenesis and highlight the importance of considering autoimmune potential when developing therapeutic interventions to reduce adverse reactions.

## Introduction

The coronavirus SARS-CoV-2 is the causative agent of the COVID-19 pandemic. COVID- 19 is an infectious disease whose typical symptoms include fever, cough, shortness of breath (Guan et al., 2020; Wang et al., 2020), and loss of taste or smell (Dawson et al., 2021). Curiously, despite hundreds of millions of confirmed cases worldwide, roughly one third are estimated to be asymptomatic (Sah et al., 2021). Yet, other SARS-CoV-2 infected individuals may also experience a variety of disease-related complications including liver injury (Saviano et al., 2021), kidney injury (Han and Ye, 2021), and cardiovascular complications including myocarditis, heart failure, thrombosis (Long et al., 2020), and thrombocytopenia (Mei et al., 2020). COVID-19 can trigger a range of antibody response levels (Wei et al., 2021) and an enrichment in autoantibodies that react with human proteins has been found for patients with severe disease (Wang et al., 2021). While the reason for the variety of disease severity affecting people with COVID-19 is not well understood, molecular mimicry may provide an avenue for explanations.

Molecular mimicry occurs when unrelated proteins share regions of high molecular similarity, such that they can perform similar and unexpected interactions with other proteins. When molecular mimicry involves antigens to which antibodies are produced, cross-reactive antibodies can result. Molecular mimicry between pathogen antigens and human proteins can cause an autoimmune response, where antibodies against the pathogen erroneously interact with human proteins, sometimes leading to transient or chronic autoimmune disorders (Getts et al., 2013). Alternatively, molecular mimicry could be viewed through the lens of heterologous immunity, where previous exposure to one pathogen antigen can result in short- or long-term complete or partial immunity to another pathogen with a similar antigen (Agrawal, 2019). Moreover, antigen-driven molecular mimicry can also lead to cross-reactive antibody immunity which has been reported against SARS-CoV-2 for uninfected individuals (Fraley et al., 2021).

The SARS-CoV-2 Spike protein is responsible for enabling SARS-CoV-2 entry into host cells (Shang et al., 2020). Spike protrudes from the virus surface and is one of the main antigenic proteins for this virus (Voss et al., 2021). Additionally, Spike is the primary component in the vaccines against SARS-CoV-2. Consequently, mimicry between Spike and human proteins or Spike and other human pathogens can result in cross-reactive antibodies in response to SARS-CoV-2 infection or vaccination. Cross-reactive antibodies may yield complex outcomes such as diverse symptoms with varying severity across populations and developmental stages as observed for COVID-19. Identifying autoimmune potential and heterologous immunity through instances of molecular mimicry between Spike and proteins from humans or human pathogens can serve to better understand disease pathogenesis, improve therapeutic treatments, and inform vaccine design as they relate to SARS-CoV-2 infection. We set out to investigate molecular mimicry between Spike and known epitopes from the Immune Epitope Database (IEDB) (Vita et al., 2019). We define molecular mimicry as a match of at least 5 identical consecutive amino acids that appear in both Spike and in a known epitope, where at least 3 amino acids are surface accessible on Spike and the match from the epitope has high structural similarity to the corresponding sequence from Spike. In light of our findings, we discuss autoimmune potential and heterologous immunity with implications for vaccine design and the side-effects of SARS-CoV-2 infection.

## Results and Discussion

We used Epitopedia (Balbin et al., 2021) to predict molecular mimicry for the structure of the SARS-CoV-2 Spike protein (PDB id: 6XR8, chain A (Cai et al., 2020)) against all linear epitopes in IEDB, excluding those from Coronaviruses. Epitopedia returned 789 sequence-based molecular mimics (termed as “1D-mimics”). 1D-mimics are protein regions from epitopes that share at least five consecutive amino acids with 100% sequence identity to a pentapeptide in SARS-CoV-2 Spike, where at least three of the amino acids are surface accessible on Spike. Most 1D-mimics (627 epitopes) were found in human. Additionally, 1D-mimics were found in non-human vertebrates (65 epitopes, 7 species), viruses (58 epitopes, 17 species), bacteria (18 epitopes, 7 species), parasitic protists (12 epitopes, 2 species), plants (5 epitopes, 2 species), and invertebrates (4 epitopes, 2 species). Seemingly redundant 1D-mimics from the same protein may represent different epitopes and, thus, all 789 1D-mimics were kept at this step.

Structural representatives from the Protein Data Bank (PDB) were identified for 284 1D- mimics based on their source sequence using the minimum cutoffs of 90% for sequence identity and 20% for query cover. The 284 1D-mimics are represented by 7,992 redundant structures from 1,514 unique PDB chains. From these, structure-based molecular mimics (termed as “3D- mimics”) were identified. 3D-mimics are 1D-mimics that share structural similarity with SARS-CoV-2 Spike as determined by an RMSD of at most 1 Å. We found 20 3D-mimics for Spike. Unsurprisingly, as with the 1D-mimics, most 3D-mimics were found for human proteins. Additionally, one 3D-mimic was found for *Mus musculus* (mouse), *Mycobacterium tuberculosis*, *Phleum pratense* (Timothy grass), and respiratory syncytial virus, respectively (Table 1). For each 3D-mimic, Epitopedia computes a Z-score based on the distribution of RMSD values for all resulting hits for the input structure. This allows for a comparative assessment of the quality of a hit, with respect to RMSD, to other hits for a given run. Epitopedia also computes an EpiScore for each hit. EpiScore, calculated as (*motif length* / *RMSD*), favors longer motifs over shorter ones with the same RMSD values.

**Table 1.**
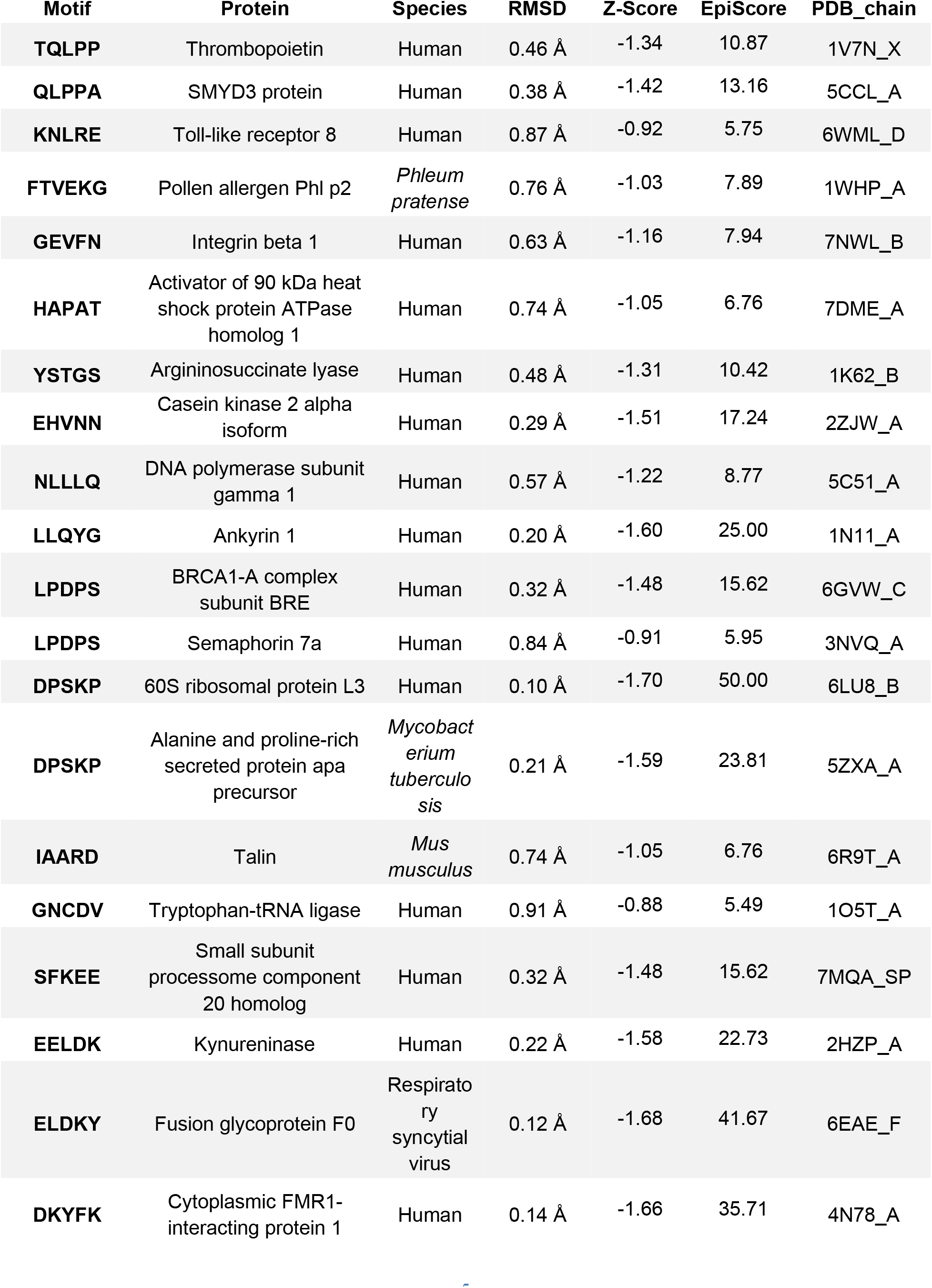
3D-mimics found for SARS-CoV-2 Spike

| Motif | Protein | Species | RMSD | Z-Score | EpiScore | PDB_chain |
| --- | --- | --- | --- | --- | --- | --- |
| **TQLPP** | Thrombopoietin | Human | 0.46 Å | -1.34 | 10.87 | 1V7N_X |
| **QLPPA** | SMYD3 protein | Human | 0.38 Å | -1.42 | 13.16 | 5CCL_A |
| **KNLRE** | Toll-like receptor 8 | Human | 0.87 Å | -0.92 | 5.75 | 6WML_D |
| **FTVEKG** | Pollen allergen Phl p2 | *Phleum pratense* | 0.76 Å | -1.03 | 7.89 | 1WHP_A |
| **GEVFN** | Integrin beta 1 | Human | 0.63 Å | -1.16 | 7.94 | 7NWL_B |
| **HAPAT** | Activator of 90 kDa heat shock protein ATPase homolog 1 | Human | 0.74 Å | -1.05 | 6.76 | 7DME_A |
| **YSTGS** | Argininosuccinate lyase | Human | 0.48 Å | -1.31 | 10.42 | 1K62_B |
| **EHVNN** | Casein kinase 2 alpha isoform | Human | 0.29 Å | -1.51 | 17.24 | 2ZJW_A |
| **NLLLQ** | DNA polymerase subunit gamma 1 | Human | 0.57 Å | -1.22 | 8.77 | 5C51_A |
| **LLQYG** | Ankyrin 1 | Human | 0.20 Å | -1.60 | 25.00 | 1N11_A |
| **LPDPS** | BRCA1-A complex subunit BRE | Human | 0.32 Å | -1.48 | 15.62 | 6GVW_C |
| **LPDPS** | Semaphorin 7a | Human | 0.84 Å | -0.91 | 5.95 | 3NVQ_A |
| **DPSKP** | 60S ribosomal protein L3 | Human | 0.10 Å | -1.70 | 50.00 | 6LU8_B |
| **DPSKP** | Alanine and proline-rich secreted protein apa precursor | *Mycobacterium tuberculosis* | 0.21 Å | -1.59 | 23.81 | 5ZXA_A |
| **IAARD** | Talin | *Mus musculus* | 0.74 Å | -1.05 | 6.76 | 6R9T_A |
| **GNCDV** | Tryptophan-tRNA ligase | Human | 0.91 Å | -0.88 | 5.49 | 1O5T_A |
| **SFKEE** | Small subunit processome component 20 homolog | Human | 0.32 Å | -1.48 | 15.62 | 7MQA_SP |
| **EELDK** | Kynureninase | Human | 0.22 Å | -1.58 | 22.73 | 2HZP_A |
| **ELDKY** | Fusion glycoprotein F0 | Respiratory syncytial virus | 0.12 Å | -1.68 | 41.67 | 6EAE_F |
| **DKYFK** | Cytoplasmic FMR1-interacting protein 1 | Human | 0.14 Å | -1.66 | 35.71 | 4N78_A |

For the 402 human 1D-mimics where no PDB structural representative could be identified for their source sequence, AlphaFold2 3D models were used. 3D model representatives were found for 102 human 1D-mimics. Of these, 10 are predicted to be AF-3D-mimics based on the RMSD (Table 2).

**Table 2.** Human AF-3D-mimics for SARS-CoV-2 Spike

| Motif | Protein | RMSD | Z-Score | EpiScore | AlphaFold2 ID |
| --- | --- | --- | --- | --- | --- |
| **NLLLQ** | Ankyrin 3 | 0.61 Å | -1.18 | 8.20 | AF-Q12955-F1-model_v1_A |
| **LLQYG** | Olfactory receptor 10Q1 | 0.66 Å | -1.13 | 7.58 | AF-Q8NGQ4-F1-model_v1_A |
| **TGIAV** | Phosphofructokinase | 0.17 Å | -1.63 | 29.41 | AF-P17858-F1-model_v1_A |
| **TGIAV** | Low affinity immunoglobulin gamma Fc region receptor II-b | 0.17 Å | -1.63 | 29.41 | AF-P31995-F1-model_v1_A |
| **KIQDSL** | Phosphorylase b kinase regulatory subunit beta | 0.19 Å | -1.61 | 31.58 | AF-Q93100-F1-model_v1_A |
| **KIQDSL** | Long-chain-fatty-acid-CoA ligase 5 | 0.37 Å | -1.43 | 16.22 | AF-Q9ULC5-F1-model_v1_A |
| **VYDPL** | Actin-binding protein IPP | 0.17 Å | -1.63 | 29.41 | AF-Q9Y573-F1-model_v1_A |
| **EELDK** | Tight junction-associated protein 1 | 0.20 Å | -1.60 | 25.00 | AF-Q5JTD0-F1-model_v1_A |
| **EELDKY** | Keratin, type I cytoskeletal 18 | 0.22 Å | -1.58 | 27.27 | AF-P05783-F1-model_v1_A |
| **ELDKY** | Tropomyosin alpha-3 chain | 0.18 Å | -1.62 | 27.78 | AF-P06753-F1-model_v1_A |

The 3D- and AF-3D-mimics (hereinafter referred to as “molecular mimics”) mapped to a few clusters on Spike. Ten molecular mimics were singletons, six overlapping molecular mimics were found as pairs in three small clusters, and the remaining 14 were found in three larger clusters with at least four overlapping molecular mimics (Figure 1a). The largest cluster, with six molecular mimics, was also adjacent to three additional molecular mimics. All mimics are displayed on the surface of Spike’s functional trimer, but the large cluster centered around LLLQY is in a deep pocket and is an unlikely antibody binding epitope in this conformation (Figure 1b). To further evaluate the autoimmune potential of the human mimics, we identified all occurrences of the motifs in the human RefSeq Select proteome (“NCBI RefSeq Select,” n.d.). The pentapeptides from the molecular mimicry regions are found from four to 33 times in human proteins (Figure 1c, Table S1). The human protein thrombopoietin that includes the 3D-mimic TQLPP (Table 2) also has an occurrence of the sequence mimic LPDPS (Table S1). Further, another protein family that occurs twice for the same pentapeptide is tropomyosin. Tropomyosin alpha-3 is an AF-3D-mimic (Table 3), and tropomyosin alpha-1 has one occurrence of the same pentapeptide (ELDKY). The same motif, ELDKY, is a 3D-mimic in the fusion F0 glycoprotein of respiratory syncytial virus (Table 2). Altogether, based on the known epitopes in IEDB, heterologous immunity is rare with Spike while the autoimmune potential resembles hotspots.

**Figure 1.**
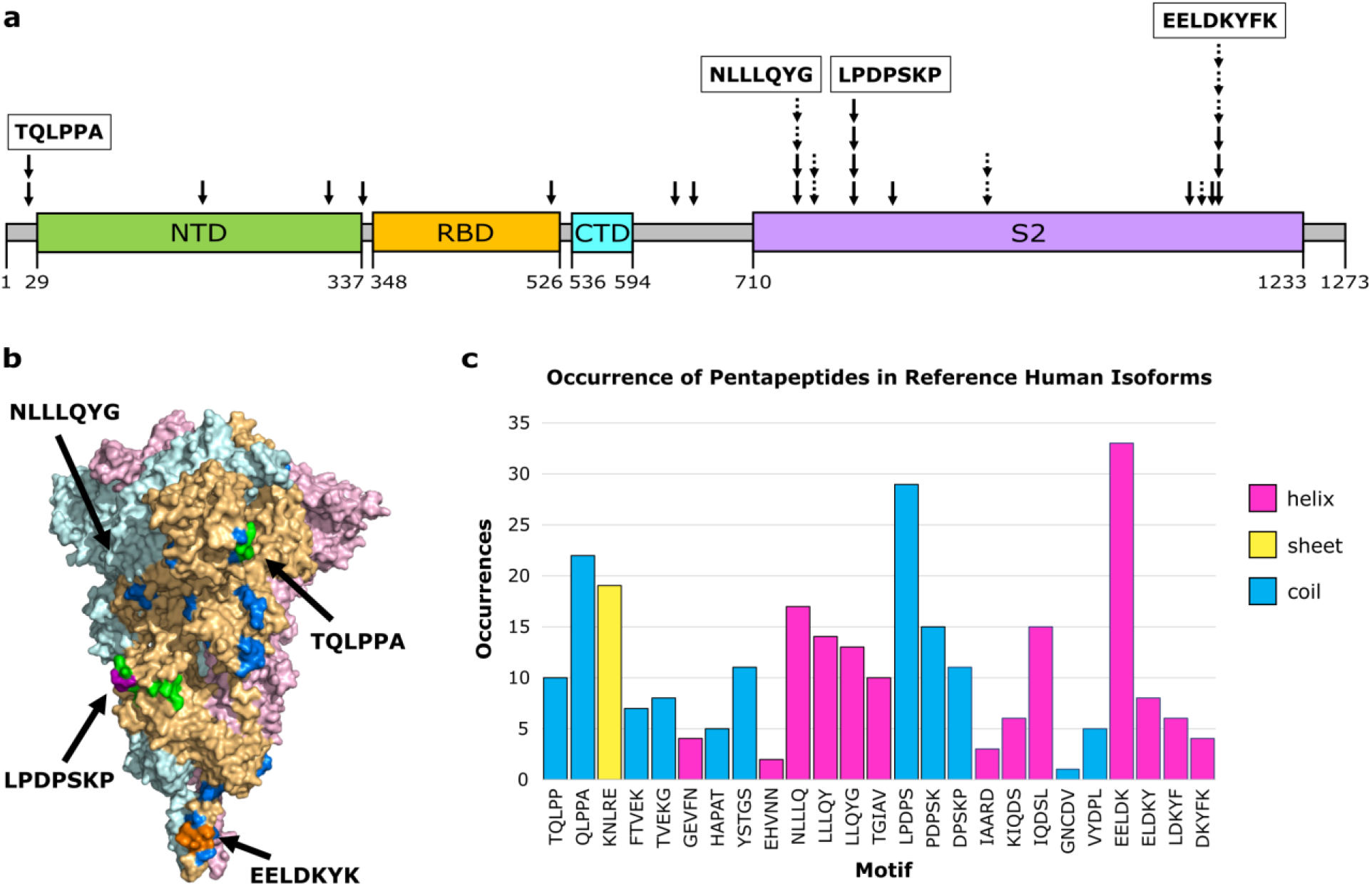
Molecular mimicry with autoimmune potential across SARS-CoV-2 Spike. (**a**) Overview of molecular mimics (solid arrow: 3D-mimic, dashed arrow: AF-3D-mimic) for Spike in the linear sequence showing Spike domains (NTD: N-terminus domain of S1 subunit, RDB: receptor binding domain of S1 subunit, CTD: C-terminus domain of S1 subunit, S2: S2 domain) as predicted by Pfam (Finn et al., 2014) based on the NCBI reference sequence (YP:009724390.1). (**b**) Surface representation of Spike (PDB id: 6XR8 (Cai et al., 2020)) colored by subunit with residues colored by number of occurrences in a molecular mimic (blue: 1, green: 2, purple: 3, orange: 4 or more). Structural visualization generated with PyMOL (Schrödinger, 2015). (**c**) The number of occurrences of the sequence motif in human RefSeq Select isoforms arranged in order from the N-terminus to the C-terminus and colored by primary secondary structure element based on Spike PDB id 6XR8.

To further evaluate molecular mimicry and, indirectly, autoimmune potential, we performed a deeper investigation of two motifs, TQLPP and ELDKY, that mapped to positions 22-26 (small cluster) and 1151-1155 (largest cluster) in Spike, respectively. For TQLPP, a 3D-mimic with human thrombopoietin was identified. The only structure in our dataset where a 3D-mimic was located at an antibody interface was for human thrombopoietin (hTPO). Thrombopoietin is a cytokine that regulates platelet production (Varghese et al., 2017) (Figure S1). Interestingly, COVID-19 patients often suffer from thrombocytopenia, characterized by low platelet levels (Yang et al., 2020), which correlates with an almost 5-fold increase in mortality (Shi et al., 2021). Thrombocytopenia in COVID-19 patients resembles immune thrombocytopenia, where hTPO and/or its receptor are mistakenly targeted by autoantibodies leading to reduced platelet count (Nazy et al., 2018). Treatments with hTPO Receptor Agonists improve thrombocytopenia in both general (Audia and Bonnotte, 2021) and COVID-19 (Watts et al., 2021) patients, suggesting the mistaken targeting occurs before hTPO activates the hTPO receptor. For ELDKY, we identified one 3D-mimic in the fusion F0 glycoprotein of respiratory syncytial virus (Table 2) and two AF- 3D-mimics from keratin type I cytoskeletal 18 and tropomyosin alpha-3 (Table 3). Additional 3D- mimics partially overlapping with ELDKY were identified. The ELDKY motif in Spike is found in an α-helix located towards the C-terminus. This motif is conserved across beta-coronaviruses and can bind an antibody effective against all human-infecting beta-coronaviruses (Pinto et al., 2021). Altogether, the numerous molecular mimics of the ELDKY motif suggests a potential for both protective and autoimmune cross-reactivity.

### Molecular mimicry between Spike and Thrombopoietin mediated through TQLPP

The shared five-amino acid motif, TQLPP (Figure 2a), is located on the surface of Spike’s N-terminal Domain (NTD) (Figure 2b, c), whereas it is found at the interface with a neutralizing antibody in hTPO (Feese et al., 2004) (Figure 2d). The TQLPP motifs from the two proteins are found in coil conformations but exhibit high structural similarity (Figure 2e, f). On Spike, the motif is adjacent to the NTD supersite that is known to be targeted by neutralizing antibodies (Cerutti et al., 2021). We hypothesized that COVID-19 may trigger the production of TQLPP-specific antibodies against this epitope that can cross-react with hTPO. In the absence of Spike TQLPP antibodies, we used molecular modeling and machine learning to investigate the binding of the neutralizing mouse Fab antibody (TN1) from the hTPO structure (Tahara et al., 1998) to the Spike TQLPP epitope.

**Figure 2.**
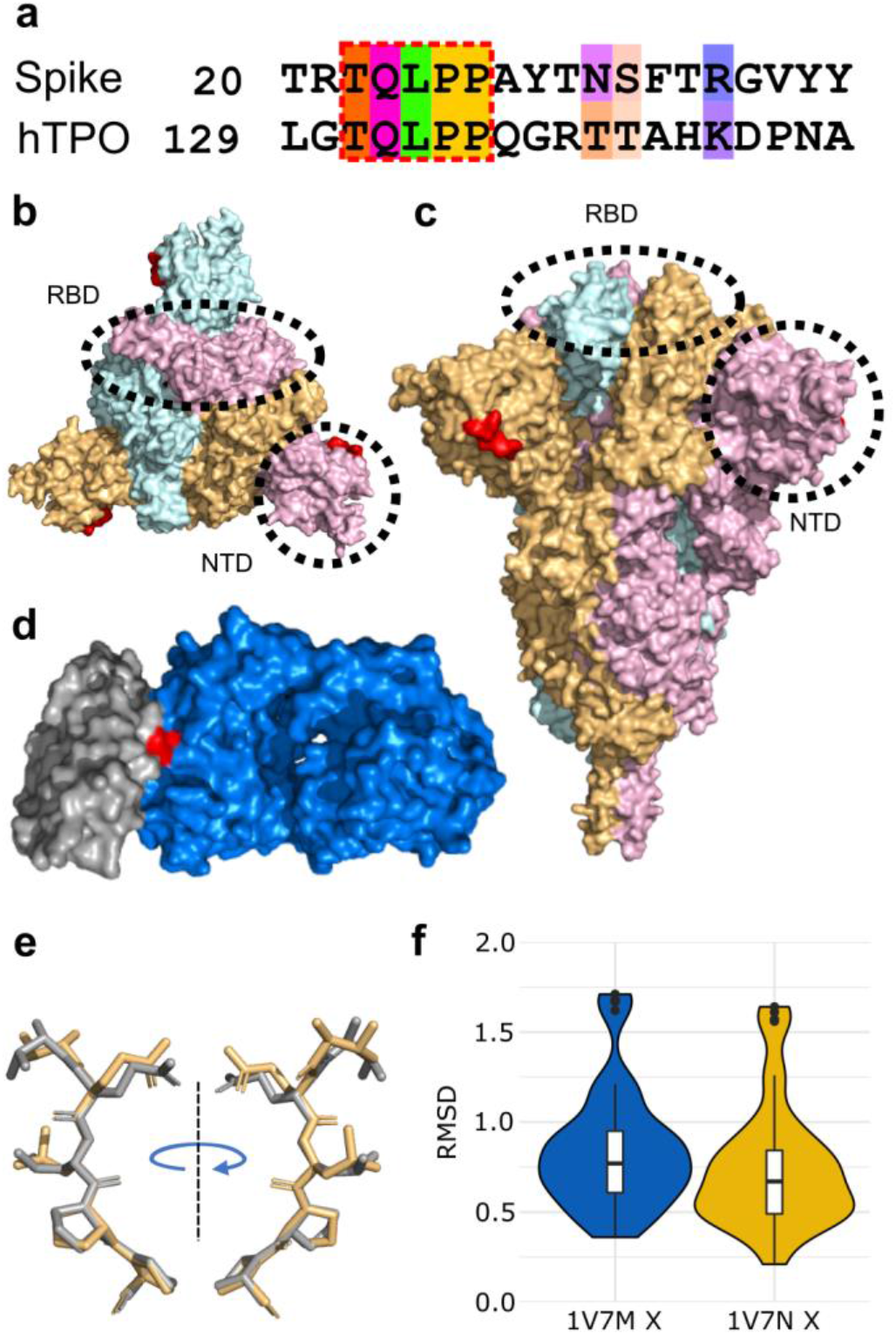
Structural mimicry between a TQLPP motif in SARS-CoV-2 Spike and an antibody binding epitope in thrombopoietin. (**a**) Pairwise sequence alignment for the TQLPP motif in the epitope for human thrombopoietin (hTPO, IEDB Epitope ID: 920946) and Spike, colored by Taylor (Taylor, 1997) for sites with ≥ 50% conservation in the amino acid property (Waterhouse et al., 2009). The region of molecular mimicry is highlighted in the red dashed box. Surface representation of Spike from (**b**) the top and (**c**) the side, with Spike trimer (PDB id: 6XR8 (Cai et al., 2020)) colored by subunit and red indicating the location of the TQLPP epitope fragment, illustrating the surface accessibility of TQLPP. (**d**) Surface representation shown for hTPO (gray, PDB id: 1V7M (Feese et al., 2004)) and its TN1 antibody (blue) with the TQLPP motif (red) at the interface. (**e**) TM-align generated structural alignment for TQLPP in Spike (beige) and hTPO (gray), with RMSD = 0.61 Å. (**f**) Violin plots of RMSD values resulting from the comparison of the TQLPP region in 60 Spike structures vs TQLPP in two hTPO structures (PDB ids: 1V7M and 1V7N, chain X for both (Feese et al., 2004)). Statistical analysis with Mann-Whitney U reveals no statistical significance between the sets. Box plots, bounded by the 1st and 3rd quartiles, show median value (horizontal solid bold line), vertical lines (whiskers) represent 1.5 × IQR, while outliers are marked as black points. For further details, see methods. Alignment representations were generated with Jalview (Waterhouse et al., 2009) and structural visualizations were generated with PyMOL (Schrödinger, 2015).

To construct a composite model of Spike and TN1 Fab, a full-length glycosylated model of the Spike trimer, based on PDB id 6VSB (Wrapp et al., 2020) with the first 26 residues (including the TQLPP motif) reconstructed (Woo et al., 2020), was coupled to three copies of TN1 Fab from the structure of hTPO complexed with TN1 Fab (Feese et al., 2004). The Spike-TN1 complex was energy minimized and equilibrated with molecular dynamics (MD) simulation. The final model of the Spike trimer complexed with three TN1 Fab antibodies (Figure 3a, b) shows that the TQLPP epitope is accessible to the antibody and the adjacent glycan chains do not shield the antibody- binding site (Figure 3c, Figure S2). To confirm the conformation of TQLPP, we calculated the RMSD for TQLPP regions from 60 Spike proteins from PDB, plus the modeled states (before and after equilibration, and upon 200 ns MD simulation) in an all-vs-all manner (Figure S3). The reconstructed TQLPP region falls within the conformational ensemble from PDB, suggesting that the modeled representation of TQLPP is valid.

**Figure 3.**
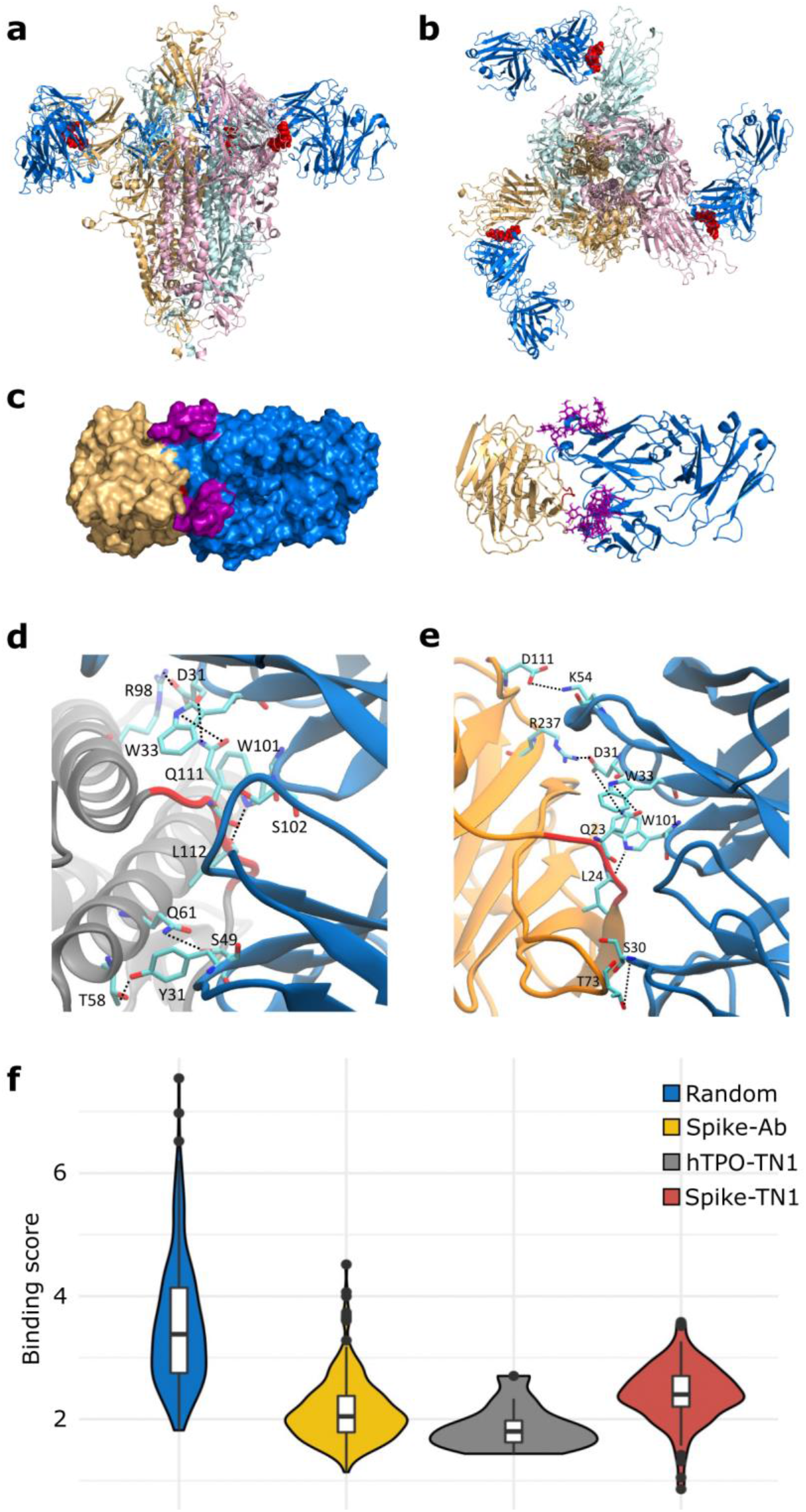
Binding of SARS-CoV-2 Spike to TN1 Fab antibody. Equilibrated structure (1 ns) of the modeled TN1 Fab antibody (blue, PDB id: 1V7M) complexed with Spike trimer model shown from (**a**) the side and (**b**) the top, with TQLPP shown as red spheres. (**c**) The Spike NTD (beige) and TN1 Fab complex used for MD simulations (200 ns), with adjacent glycans at N17 and N74 highlighted in purple. The representative amino acids contributing to hydrogen bonds (dashed lines) during the last 50 ns of simulations for the (**d**) hTPO-TN1 and (**e**) Spike-TN1 complexes are highlighted as cyan sticks. (**f**) Violin plot showing the distribution of the MaSIF binding score values for randomly selected patch pairs (blue), the interacting region of Spike-antibody (yellow) and hTPO-TN1 (gray) complexes, and for modeled Spike-TN1 complexes across 40 Spike configurations (red). Statistical analysis with Mann-Whitney U shows that all pairwise comparisons except for Spike-Ab and hTPO-TN1 are significantly different after Bonferroni correction (Table S5). Box plots, bounded by the 1st and 3rd quartiles, show median value (horizontal solid bold line), vertical lines (whiskers) represent 1.5 × IQR, while outliers are marked as black points. For further details, see methods. Structural visualizations were generated with PyMOL (Schrödinger, 2015) and VMD (Humphrey et al., 1996).

To evaluate the molecular mimicry between the antibody interface areas, we performed MD simulations of hTPO and Spike NTD with TQLPP complexed with the TN1 antibody. The hydrogen bonds were calculated between the TN1 antibody with hTPO and Spike, respectively, from the last 50 ns of both trajectories (Figure S4). Both the Spike-TN1 and the hTPO-TN1 complexes showed similar contact areas (Figure S4). Notably, critical hydrogen bonds were observed for residues Q and L in the TQLPP motif with TN1 for both Spike and hTPO (Figure 3d, e and Figure S4).

To further support our findings, we evaluated the antibody-antigen interface complementarity with MaSIF-search, a recent tool that uses deep learning techniques (Gainza et al., 2019), on a pair of circular surface regions (patches) from an antibody-antigen complex. MaSIF-search produces a score associated with the strength of binding when forming a stable complex. Lower scores are associated with stronger binding. We refer to this score here as the binding score. The results show that Spike-TN1 complexes have a better (lower) binding score than random complexes and that complexes including Spike from PDB ID 7LQV (Cerutti et al., 2021) have three of the four best binding scores (0.86, 1.05, 1.42) and may bind to TN1 as well as, or better than, hTPO (Figure 3h, Tables S4-5). Notably, in 7LQV, COVID-19 antibodies bind to Spike at the NTD supersite (Cerutti et al., 2021). These results strongly argue for the possibility of cross-reactivity between Spike and hTPO driven by the molecular mimicry of TQLPP (Figure 3).

The human proteome contains nine additional occurrences of the TQLPP motif. Two of these motifs, found in Hermansky-Pudlak syndrome 4 protein and ALG12 (Mannosyltransferase ALG12 homolog), have been associated with thrombosis and hemostasis disorder (Kanduc, 2021). To evaluate structural mimicry between Spike-TQLPP and all human-TQLPP motifs, we utilized AlphaFold2 3D models (Jumper et al., 2021; Tunyasuvunakool et al., 2021) (Figure S5). The closest structural mimicry region is in hTPO (RMSD = 0.39 Å), followed by coiled-coil domain-containing protein 185, Fc receptor-like protein 4 (FCRL4), and far upstream element-binding protein 1 (Figure S5). These results indicate that TQLPP motifs have similar conformations (Figure S3), strengthening the notion of structural mimicry. We investigated the potential cross-reactivity of an antibody targeting TQLPP in these proteins, after discarding six that display the TQLPP motif in low confidence or unstructured regions. The remaining three proteins, NEK10 (ciliated cell-specific kinase), FCRL4, and ALG12 were complexed with TN1 (Figure 4). The binding score for NEK10-TN1 (1.44) is comparable to the hTPO-TN1 complex (Figure 4). NEK10 regulates motile ciliary function responsible for expelling pathogens from the respiratory tract (Chivukula et al., 2020). Dysfunction of NEK10 can impact mucociliary clearance and lead to respiratory disorders such as bronchiectasis (Chivukula et al., 2020). Based on our results, it is plausible that the function of NEK10 and thus mucociliary clearance can be affected by cross-reactive Spike antibodies targeting TQLPP.

**Figure 4.**
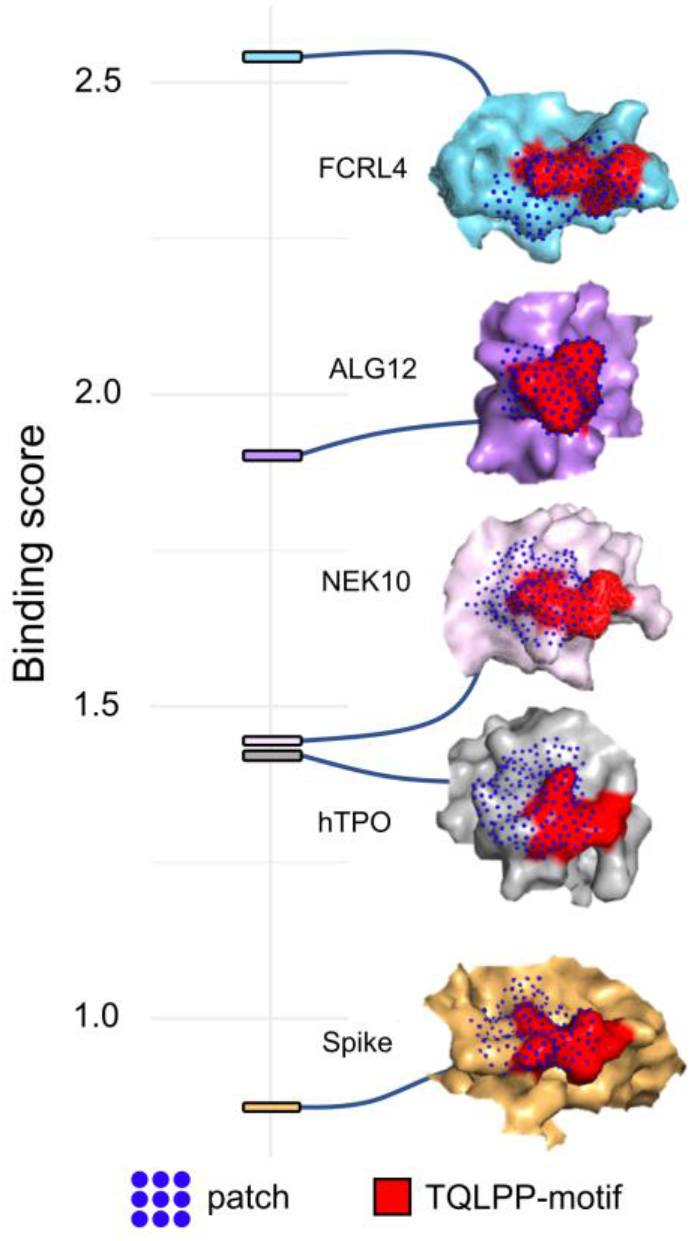
Predicted interaction patches between TN1 Fab antibody (PDB id: 1V7N) and the TQLPP motif. The best (lowest) binding score is shown for Spike (PDB id: 7LQV, chain A), hTPO (PDB id: 1V7N, chain X), NEK10 (Uniprot: Q6ZWH5), ALG12 (Uniprot: Q9BV10), and FCRL4 (Uniprot: Q96PJ5). For all, red indicates the TQLPP motif and blue dots represent the surface points included in the predicted MaSIF patches.

### Molecular mimicry between Spike, RSV, and many human proteins mediated through ELDKY

Another motif, ELDKY, is in a region with several partially overlapping pentamer motifs including three 3D-mimics and three AF-3D-mimics (Figure 5a). For the 3D-mimics, two are from the human proteins kynureninase (hKYNU; motif: EELDK) and cytoplasmic FMR1-interacting protein 1 (hCYFIP1; motif: DKYFK), while the last is found in the fusion F0 glycoprotein of respiratory syncytial virus (RSV; motif: ELDKY). For the AF-3D-mimics, the motif is found in human tight junction-associated protein 1 (hTJAP1; motif: EELDK), keratin type I cytoskeletal 18 (hkRT18; motif: EELDKY), and tropomyosin alpha-3 (hTPM3; motif: ELDKY). In Spike, the ELDKY motif is in a stem helix region near the C-terminus. This motif is well-conserved across beta- coronaviruses and has been shown to bind to a broadly neutralizing antibody effective against all human-infecting beta-coronaviruses (Pinto et al., 2021). Additionally, stronger antibody responses to the epitope containing the ELDKY motif have been recorded for severe (requiring hospitalization) vs moderate cases, while fatal cases had a weaker response than surviving cases (Voss et al., 2021). Together with the 3D-mimics identified here, these results suggest interesting possibilities for the ELDKY motif from the perspective of both protective immunity and an autoimmune response. First, while not an example of molecular mimicry, prior exposure to an endemic cold-causing coronavirus (ex. HCoV-OC43) could result in the production of a broadly neutralizing antibody against an epitope containing the ELDKY motif that would be effective against SARS-CoV-2 infection, which could result in milder or asymptomatic infection. Further, a protective effect due to molecular mimicry is suggested by the 3D-mimic identified for the fusion F0 glycoprotein of RSV, a common virus that infects most children in the United States by the time they are 2 years old (“Respiratory Syncytial Virus (RSV) | NIH: National Institute of Allergy and Infectious Diseases,” n.d.), where antibodies against the ELDKY-containing epitope in RSV may be effective in combatting SARS-CoV-2 infection. In contrast, the potential for an autoimmune response against this motif is suggested by its presence in both two human 3D- and AF-3D-mimics (Figure 5).

**Figure 5.**
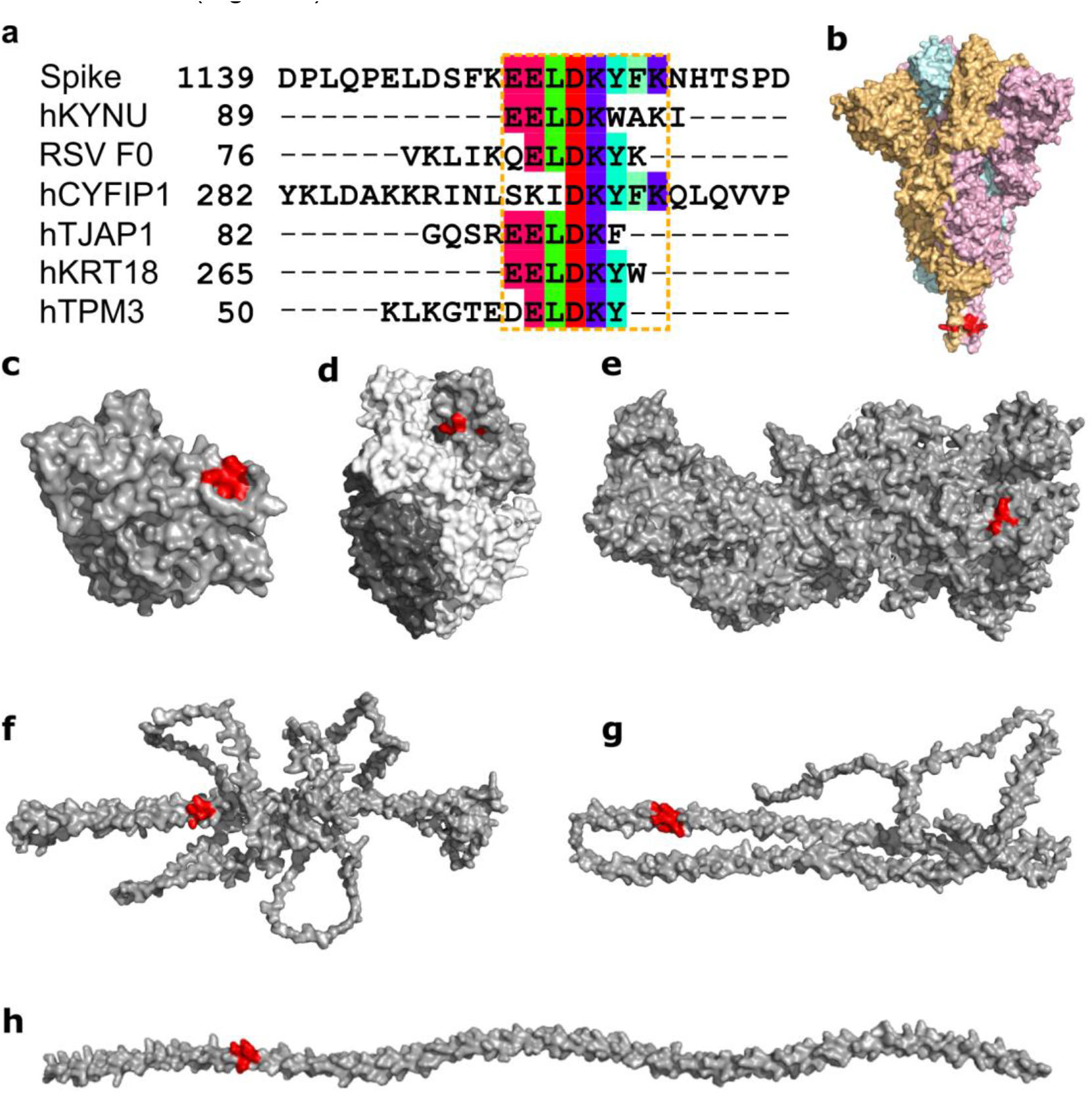
Structural mimicry between an ELDKY motif in SARS-CoV-2 Spike and epitopes in 6 other proteins. (**a**) Sequence alignment between SARS-CoV-2 Spike and the epitopes containing the 3D-mimicry motif for human kynureninase (hKYNU, IEDB Epitope ID: 1007556), respiratory syncytial virus fusion F0 glycoprotein (RSV F0, IEDB Epitope ID: 1087776), human cytoplasmic FMR1-interacting protein 1 (hCYFIP1, IEDB Epitope ID: 1346528), human tight junction-associated protein 1 (hTJAP1, IEDB Epitope ID: 1016424), human keratin type I cytoskeletal 18 (hKRT18, IEDB Epitope ID: 1331545), and human tropomyosin alpha-3 (hTPM3, IEDB Epitope ID: 938472). Residues in the molecular mimicry motifs are colored by Taylor (Taylor, 1997). The extended molecular mimicry region is highlighted by the orange dashed box. (**b**) Surface representation of Spike (PDB id: 6XR8) colored by subunit with ELDKY motif indicated in red. Surface representation of proteins (gray) with full or partial 3D-mimics of the ELDKY motif (red): (**c**) hKYNU (PDB id: 2HZP), (**d**) RSV F0 (PDB id: 6EAE), (**e**) hCYFIP1 (PDB id: 4N78), (**f**) hTJAP1 (Uniprot: Q5JTD0), (**g**) hKRT18 (Uniprot: P05783), (**h**) hTPM3 (Uniprot: P06753). Alignment representations were generated with Jalview (Waterhouse et al., 2009) and structural visualizations were generated with PyMOL (Schrödinger, 2015).

There are six additional occurrences of the ELDKY motif in the human proteome (Figure S6). Structural similarity between Spike-ELDKY and human-ELDKY was assessed based on experimentally determined structures (if available) or AlphaFold2 3D models. RMSDs for the ELDKY motif ranged from 0.12-0.20 Å for 5 of the structures, with one hit being an outlier at an RMSD = 0.46 Å. In all instances, the ELDKY motif is found in an α-helix, resulting in the high degree of structural similarity found for this motif across proteins and bolstering the possibility for molecular mimicry. The ELDKY occurrence with the largest RMSD (0.46 Å) is found in the leucine-zipper dimerization domain of cGMP-dependent protein kinase 1 (PRKG1) (Figure S6) whose phosphorylation targets have roles in the regulation of platelet activation and adhesion (Li et al., 2003), smooth muscle contraction (Sauzeau et al., 2000), and cardiac function (Francis, 2010). Additionally, PRKG1 regulates intracellular calcium levels via a multitude of signaling pathways (Francis et al., 2010). The ELDKY motif is also found in tropomyosin alpha-1 (TPM1), a homolog of the AF-3D-mimic tropomyosin alpha-3 (TPM3). Tropomyosins (TPMs) are involved in regulation of the calcium-dependent contraction of striated muscle (Szent-Györgyi, 1975). TPM1 is a 1D-mimic but due to a discrepancy in IEDB it was not identified as a 3D-mimic, although there is high structural similarity between ELDKY in Spike and ELDKY in TPM1 (Figure S6). A previous study identified a longer match with 53% sequence identity between Spike and TPM1 that included the ELDKY motif but was not able to show the structural similarity (Marrama et al., 2022) due to using a structure for Spike that did not include the ELDKY motif. Cross-reactive Spike antibodies targeting ELDKY may react with PRKG1, affecting its role in the regulation of platelet activation and adhesion and thus providing another avenue for thrombocytopenia or other blood clotting disorders. Antibodies that cross-react with PRKG1 may also alter calcium levels, thus affecting TPM function. For TPM1, cross-reactive Spike antibodies targeting the ELDKY motif may result in coronary heart disease, as low-level autoantibodies against this protein have been associated with increased risk for this condition (Zhang et al., 2020) and TPM1 and TPM3 are cardiac disease-linked antigens (Marrama et al., 2022). Cardiac disease, including myocardial injury and arrhythmia, can be induced by SARS-CoV-2 infection (Nishiga et al., 2020) and myocarditis has been found to develop in some individuals following vaccination against SARS-CoV-2 (Patone et al., 2021). Furthermore, COVID-19 has been found to increase risk and long-term burden of several cardiovascular diseases, with COVID-19 severity being proportionate to increased risk and incidence. (Xie et al., 2022).

## Conclusion

We find that molecular mimics with high autoimmune potential are often found in clusters within Spike. Some clusters have several molecular mimics whose motifs are also found multiple times in the human proteome. Molecular mimics located in α-helices tend to have high structural similarity as can be expected based on the regular conformation of the helix, but also some molecular mimics in coil regions are remarkably similar. Our results on the TQLPP motif, located in a coil region, suggest a worrisome potential for cross-reactivity due to molecular mimicry between Spike and hTPO involving the TQLPP epitope that may affect platelet production and lead to thrombocytopenia. Further, cross-reactivity with other TQLPP-containing proteins such as NEK10 cannot be dismissed based on our in-silico results, but in-vivo validation is required. The presence of neutralizing antibodies against peptides with TQLPP in COVID-19 patients’ convalescent plasma (Li et al., 2020), particularly in severe and fatal cases (Voss et al., 2021) adds credence to our result. It is also interesting to note that antibodies against a TQLPP- containing peptide were found in the serum of pre-pandemic, unexposed individuals (Stoddard et al., 2021). Prior infection with a different human coronavirus cannot explain the cross-reactivity observed in the unexposed group because TQLPP is situated in a region with low amino acid conservation (Stoddard et al., 2021). Rather, this suggests the presence of an antibody for an unknown epitope with affinity for the TQLPP region in Spike. The COVID-19 vaccines designed to deliver the Spike protein from SARS-CoV-2, like COVID-19 infection itself, can cause thrombocytopenia (Greinacher et al., 2021; Helms et al., 2021; Schultz et al., 2021; Yang et al., 2020) and it is plausible that cross-reactivity can titrate the serum concentration of free hTPO. The evolutionary trends in the TQLPP motif suggested it may not remain in Spike. In the Gamma variant, a P26S mutation has changed TQLPP to TQLPS and two additional mutations are located just before the motif at L18F and T20N in the NTD supersite, while the Delta variant is mutated at T19R (Hodcroft, 2021). The first Omicron variant (21K or BA.1), however, has no amino acid substitutions near the TQLPP motif, while a closely related Omicron variant (21L or BA.2) contains a 9 nucleotide deletion that results in the loss of 60% of the TQLPP motif (L24 -, P25-, P26-) (Hodcroft, 2021). Neutralizing antibodies targeting the NTD supersite may rapidly lose efficacy against the evolving SARS-CoV-2. Consequently, protein engineering of the TQLPP motif and possibly the NTD supersite for future COVID-19 vaccines may reduce the risk for thrombocytopenia and improve long-term vaccine protection against evolving variants.

We illuminated the cross-reactivity mediated through the ELDKY motif between Spike and PRKG1, TPM1, and TPM3. While PRKG1 provides a connection between blood clotting disorders and cardiac complications, it has a larger RMSD than other ELDKY motifs. ELDKY motifs in α- helices have high similarity and make good candidates for molecular mimicry. We find ELDKY in the homologous proteins TPM1 and TPM3 suggesting a conserved importance for structure and function. In contrast to TQLPP, the ELDKY motif is highly conserved among beta-coronaviruses (Pinto et al., 2021) and there are presently no SARS-CoV-2 variants with mutations in this region (Hodcroft, 2021). Further, while the existence of a broadly neutralizing antibody against an epitope containing ELDKY (Pinto et al., 2021) illustrates the potential of this motif as a pan-coronavirus vaccine target, the viability may be diminished by the possibility for autoimmune cross-reactivity due to this motif.

We present an extended application of Epitopedia (Balbin et al., 2021) to identify molecular mimicry between Spike and known epitopes. We do not attempt to discover all possible epitopes for Spike but focus on epitopes with positive assays from the IEDB (Vita et al., 2019). For one epitope, we find the TQLPP motif and an interacting antibody with which we perform a computational investigation into antibody binding properties of the tentative molecular mimic. The results show that the same antibody may be able to bind TQLPP-containing epitopes in different proteins and that the TQLPP motif tends to be found in similar conformations despite being in a loop or coil. For another epitope, we find the ELDKY motif with potential for protective immunity and with high structural similarity. High structural similarity can be expected for α-helical structures, and, if the sequence is identical, molecular mimicry results. Altogether, these are examples of molecular mimicry that may play a role in autoimmune or cross-reactive potential of antibodies generated by the immune system against SARS-CoV-2 Spike, but it must be noted that these results have not been experimentally verified. Still, computational investigations into the autoimmune potential of pathogens like SARS-CoV-2 are important for therapeutic intervention and when designing vaccines to avoid potential predictable autoimmune interference.

## Methods

### Identifying epitopes with molecular mimicry

To identify known epitopes with positive assays, we used Epitopedia (Balbin et al., 2021) with a Cryo-EM structure of Spike from SARS-CoV-2 (PDB id: 6XR8, chain A (Cai et al., 2020)) as input. Hits containing 5 or more consecutive residues with 100% sequence identity where at least 3 of the input residues are surface accessible are considered sequence-based molecular mimics (termed as “1D-mimics”). For all 1D-mimics with corresponding structural representation from either PDB (Berman et al., 2000) or AlphaFold2 (Tunyasuvunakool et al., 2021) 3D models of human proteins, TM-align (Zhang and Skolnick, 2005) was used to generate a structural alignment and Root Mean Square Deviation (RMSD) for all input-hit (1D-mimic) alignment pairs using only the structural regions corresponding to the hit for the source antigenic protein containing the epitope and the input. Epitopes with an RMSD ≤ 1 Å to Spike were considered structure-based molecular mimics (termed as “3D-mimics”).

### Conformational ensemble of TQLPP structural mimicry

To gather all structures of the TQLPP motif in Spike, an NCBI BLASTP search against PDB was performed with the SARS-CoV-2 Spike reference sequence as the query and a SARS-CoV-2 taxa filter. Of 75, close to full-length, hits (>88% query cover), 20 included a solved structure for the TQLPP motif. The TQLPP region of the PDB structure was extracted for all chains in the 20 structures (all were trimers, as in Spike’s biological state) resulting in a TQLPP Spike ensemble of 60 different chains from SARS-CoV-2. Each sequence in the TQLPP Spike ensemble was superimposed with chain X of the two PDB structures of human thrombopoietin (hTPO, PDB ids: 1V7M and 1V7N) to generate an RMSD value distribution for Spike’s conformational ensemble vs hTPO for the structural mimicry region (Table S2).

### Modeling Spike-TN1 complex

We constructed a composite model of the Spike-TN1 complex using the hTPO-TN1 complex (PDB id: 1V7M) as a template. For this, we first aligned the TQLPP segment of hTPO in the hTPO-TN1 complex with the TQLPP segment of the fully glycosylated model of Spike (PDB id: 6VSB (Wrapp et al., 2020)) retrieved from the CHARMM-GUI Archive (Choi et al., 2021). We then removed hTPO, leaving TN1 complexed with Spike. For the Spike-TN1 simulations, only the TN1 interacting N-terminal domain of Spike (residues 1-272) was considered. Geometrical alignments, as well as visualization, were performed with PyMOL version 2.5 (Schrödinger, 2015) and Visual Molecular Dynamics (VMD 1.9.3 (Humphrey et al., 1996)).

To confirm that the modeled Spike TQLPP region is in agreement with the TQLPP region of solved Spike structures, these regions were extracted. TM-align was used to superimpose the TQLPP regions from the different structures, including the modeled TQLPP region from the Spike- TN1 complex, and to calculate the respective RMSD values. Three states of the model were included (before and after equilibration, and after molecular dynamics (described in the following paragraph)) together with the 60 experimentally determined Spike structures in Table S2 and compared in an all-against-all manner (Figure S3, Table S3).

### Molecular dynamics simulation

A simulation system for the modeled Spike-TN1 complex was prepared using CHARMM-GUI (Brooks et al., 2009; Jo et al., 2008; Lee et al., 2016). The complex was solvated using a TIP3P water model and 0.15 M concentration of KCl and equilibrated for 1 ns at 303 K. All-atom simulations were performed with NAMD2.14 (Phillips et al., 2005) using CHARMM36m force-field. The production runs were performed under constant pressure of 1 atm, controlled by a Nose−Hoover Langevin piston (Nosé and Klein, 1983) with a piston period of 50 fs and a decay of 25 fs to control the pressure. The temperature was set to 303 K and controlled by Langevin temperature coupling with a damping coefficient of 1/ps. The Particle Mesh Ewald method (PME) (Essmann et al., 1995) was used for long-range electrostatic interactions with periodic boundary conditions and all covalent bonds with hydrogen atoms were constrained by Shake (Ryckaert et al., 1977). The contact area of the interface was calculated as (S1+S2-S12)/2, where S1 and S2 represent the solvent accessible surface areas of the antigen and antibody and S12 represents that for the complex (Figure S4). We performed MD simulations of the hTPO-TN1 complexes (PDB ids: 1V7M and 1V7N) as well as the Spike-TN1 complexes modeled from PDB ids: 1V7M and 1V7N to generate interaction matrices of protein-antibody hydrogen bonds during the last 50 ns of 200 ns MD simulation for each run.

### Antibody interface complementarity

We used the MaSIF-search geometric deep learning tool designed to uncover and learn from complementary patterns on the surfaces of interacting proteins (Gainza et al., 2019). Surface properties of proteins are captured using radial patches. A radial patch is a fixed-sized geodesic around a potential contact point on a solvent-excluded protein surface (Sanner et al., 1996). In MaSIF-search, the properties include both geometric and physicochemical properties characterizing the protein surface (Gainza et al., 2019). This tool exploits that a pair of patches from the surfaces of interacting proteins exhibit interface complementarity in terms of their geometric shape (e.g., convex regions would match with concave surfaces) and their physicochemical properties. The data structure of the patch is a grid of 80 bins with 5 angular and 16 radial coordinates and ensures that its description is rotation invariant. Each bin is associated with 5 geometric and chemical features: shape index, distance-dependent curvature, electrostatics, hydropathy, and propensity for hydrogen bonding. The model converts patches into 80-dimensional descriptor vectors, such that the Euclidian distance between interacting patches is minimized. Here, we define the binding score as a measure of distance between the descriptor vectors of the two patches. Thus, lower “MaSIF binding scores” represent better complementarity and therefore better matches. The pre-trained MaSIF-search model “sc05” with a patch radius of 12 Å was used.

Using the MaSIF protocol, we evaluated complexes of the TN1 antibody bound to Spike in the TQLPP region. The antibody-antigen patch pairs were extracted using scripts from the molecular mimicry search pipeline EMoMiS (Stebliankin et al., 2022). To accommodate multiple Spike configurations, we extracted patches from 40 SARS-CoV-2 Spike-antibody complexes from the SabDab structural antibody database (Dunbar et al., 2014). Patches centered at Q23 from Spike and W33 from TN1 were selected as representative pairs for the Spike-TN1 interaction type because this potential contact point has the most hydrogen bonds in the modeled Spike-TN1 TQLPP region. Binding scores of randomly formed complexes (Random), complexes between Spike and its native antibodies (Spike-Ab), and complexes between hTPO and TN1 (hTPO-TN1) were extracted and tabulated (Table S4). The distribution of binding scores from randomly formed complexes was obtained by pairing patches from random locations on Spike with patches from its antibodies. For native antibody-antigen Spike-Ab and hTPO-TN1 complexes, we obtained patch pairs from known interface regions using the MaSIF-search strategy for the selection of interacting patches (Gainza et al., 2019). Columns “Contact AB” and “Contact AG” in Table S4 indicate the residue used as the center of the patch from the antibody and the corresponding antigen.

### Evaluating further cross-reactivity

All 3D-mimics and AF-3D-mimics were split into pentapeptides (if mimicry motif exceeded 5 residues) which were used as queries for NCBI BLASTP searches against the RefSeq Select (“NCBI RefSeq Select,” n.d.) set of proteins from the human proteome. Results for the BLAST searches can be found in Table S1.

For the TQLPP sequence motif, 10 representative isoforms in proteins containing the complete motif were found, including hTPO. The other 9 proteins lacked a solved structure containing TQLPP. However, AlphaFold2 3D models were available for all 10 of these RefSeq Select sequences (Jumper et al., 2021; Tunyasuvunakool et al., 2021), allowing us to extract the region corresponding to TQLPP in these hits and structurally superimpose this region with Spike TQLPP (from PDB id 6XR8) with TM-align as described above.

TN1-protein complexes were generated for three of the remaining 9 proteins (Fc receptor-like protein 4 (residues 190-282), serine/threonine-protein kinase NEK10 (residues 1029-1146), ALG12 (Mannosyltransferase ALG12 homolog (residues 1-488)). The TQLPP segment in hTPO was structurally aligned with each of the TQLPP segments of the modeled proteins, after which, hTPO was removed resulting in the complex of TN1 with the modeled proteins following the methods mentioned for Spike above. The equilibrated structures of these complexes show that TN1 stays firmly with these proteins without any structural clash. Further, to evaluate the shape complementarity of these three proteins and TN1, MaSIF was used to calculate binding scores as described above (Table S6).

It should also be noted that two additional human genes (GeneIDs 8028 and 57110) also have one TQLPP motif, but not in the RefSeq Select isoforms. Since no structure or structural prediction was available for these proteins, they were excluded from further analysis.

For the ELDKY sequence motif, 6 additional representative isoforms containing the complete motif were found, in addition to the human proteins identified by Epitopedia to contain 3D-mimics of the motif. Solved structures of the ELDKY motif were available for 3 of the proteins, while the others had AlphaFold2 3D models available. In all instances, the region corresponding to the ELDKY motif was extracted and structurally superimposed with Spike ELDKY (from PDB id 6XR8) with TM-align as previously described.

### Statistical analysis

Distributions were visualized as violin plots with `ggpubr` and `ggplot2`. Following Shapiro- Wilk normality testing, statistical analysis comparing the different distributions was performed using Mann-Whitney U with *SciPy*, followed by a simplified Bonferroni correction (alpha/n comparisons) when appropriate.

## Acknowledgments

We thank Dr. Sathibalan Ponniah (Immune Analytics LLC, Columbia, MD), Dr. Charles Dimitroff (Florida International University), Dr. Sixto Leal (University of Alabama - Birmingham), and Kevin Bennett (Florida International University) for discussions.

## Contributions

J.S.-L., J.N.-C., C.A.B., G.N., P.C., K.M., T.C., A.M.M., P.B., and V.S. designed the overall method and approach. J.S.-L., G.N., P.C., A.M.M., K.M. supervised the research. J.N.-C., C.A.B., and J.S.-L. identified molecular mimicry. P.B. and P.C. performed modeling. G.N. and V.S. performed MaSIF-search. J.N.-C., C.A.B., J.S.-L., P.C, G.N., P.B., and V.S. analyzed the data. M.S., J.N.-C., C.A.B., J.S.-L, V.S., and P.B. performed visualization. G.N. and J.S.-L. performed project administration. J.S.-L. and J.N.-C. acted as lead authors. C.A.B., G.N., P.C., K.M., T.C., A.M.M., P.B., and V.S. contributed to writing the manuscript. All authors read and commented the manuscript.

## Corresponding author

Correspondence to Jessica Siltberg-Liberles

## Supplementary materials

**Figure S1.**
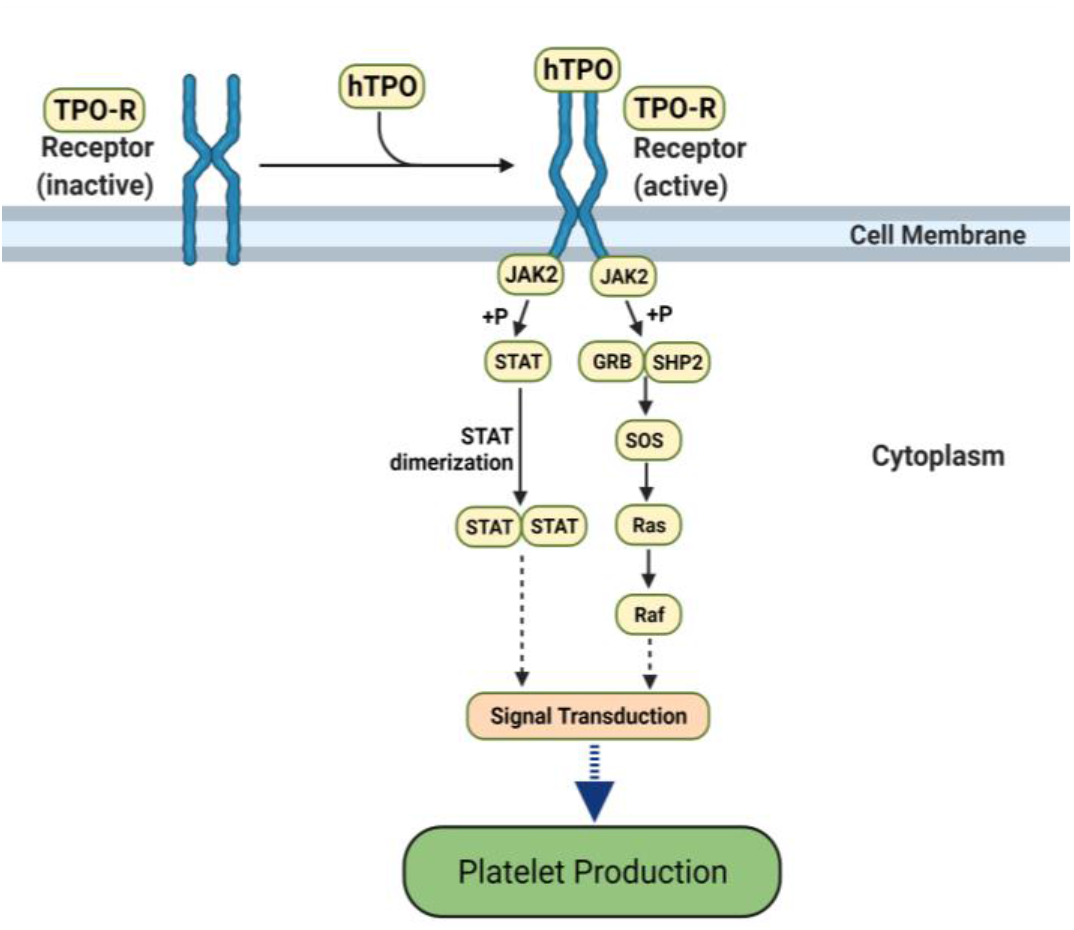
The hTPO pathway to induce platelet production. Simplified JAK-STAT signaling pathway in megakaryocytes where hTPO activates the TPO receptor and triggers signaling cascades that stimulate platelet production (Kanehisa et al., 2021; Kuter, 2013). Created with BioRender.com.

**Figure S2.**
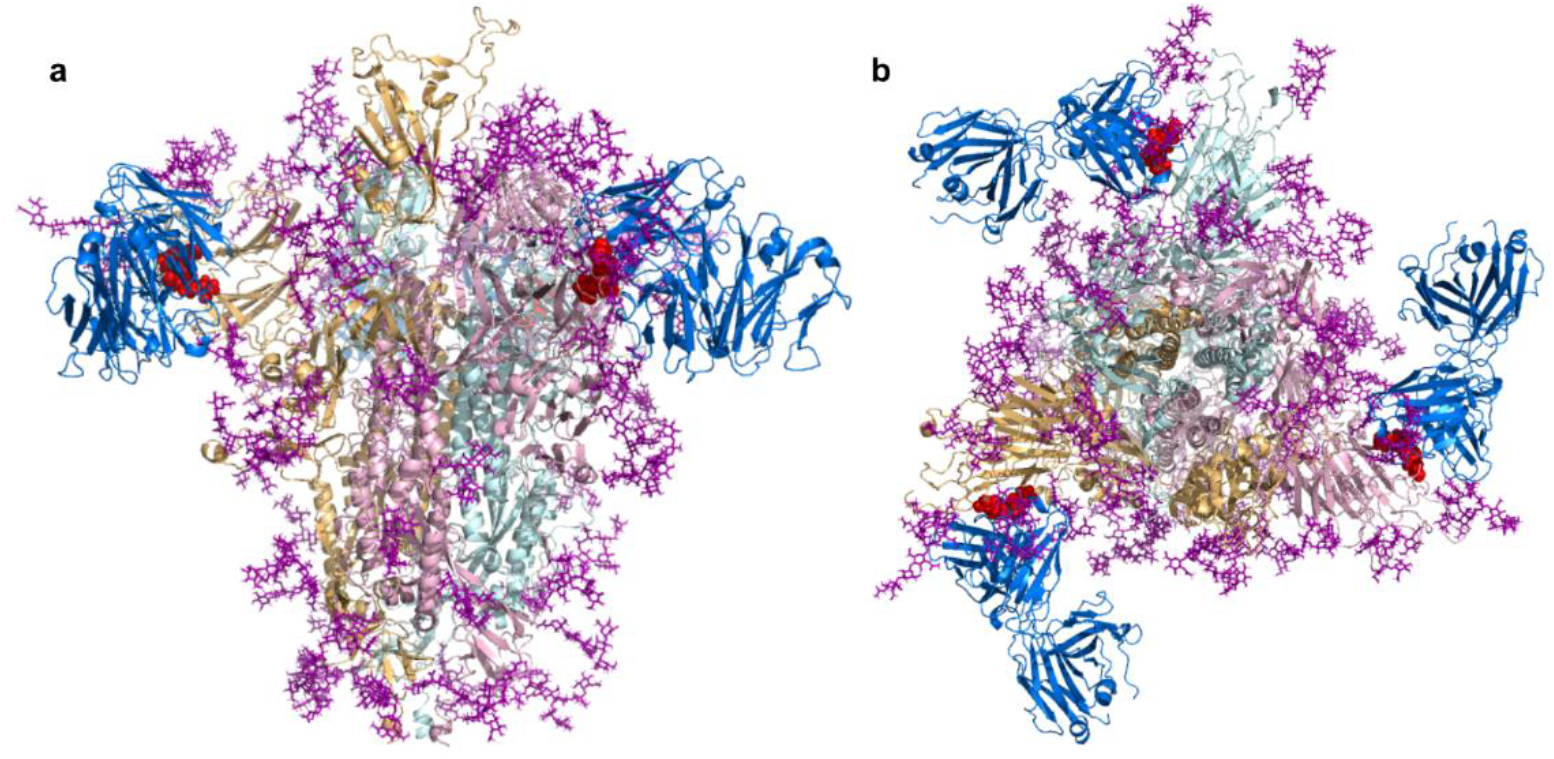
SARS-CoV-2 Spike bound to TN1 Fab antibody. SARS-CoV-2 Spike shown in the trimeric state (PDB id: 6VSB) bound to TN1 Fab antibody (blue, PDB id: 1V7M) as viewed from (**a**) the side and (**b**) the top. The TQLPP motifs are shown as red spheres and glycans are shown in purple. Structural visualization generated with PyMOL (Schrödinger, 2015).

**Figure S3.**
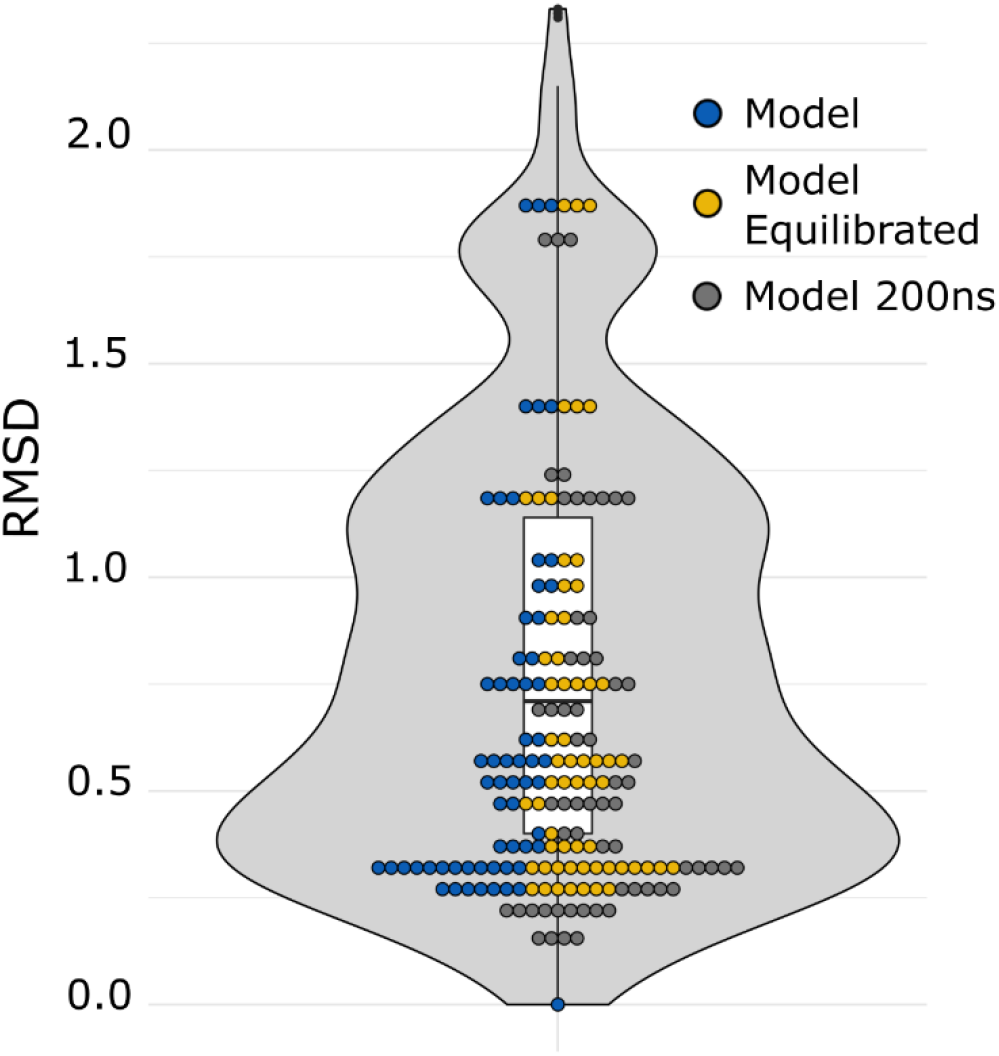
RMSD value distribution for solved and modeled Spike TQLPP regions. RMSD values resulting from an all-against-all comparison of the Spike TQLPP region of 63 structures, including the model in 3 states (shown as dots).

**Figure S4.**
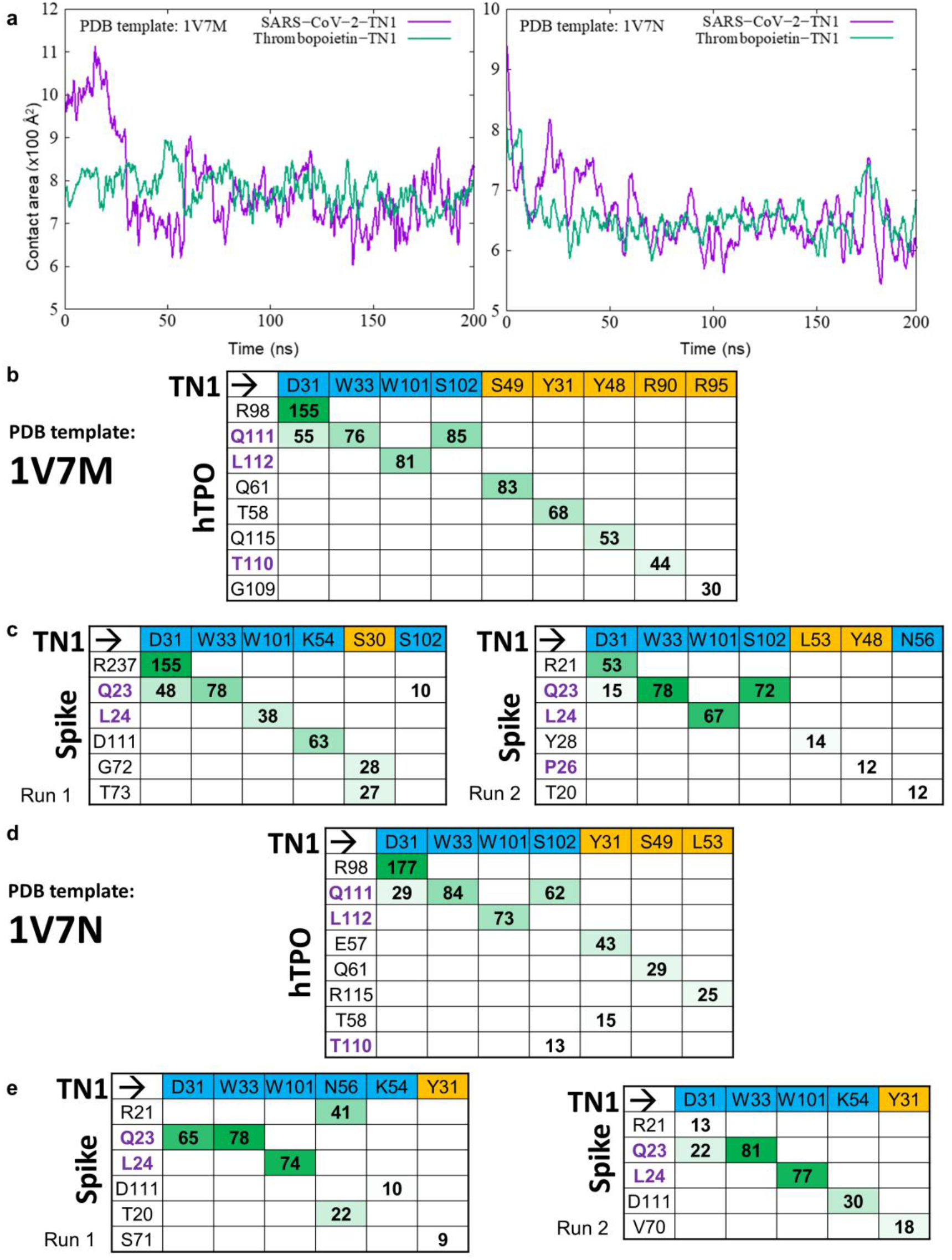
Molecular dynamics simulations overview. (**a**) Time evolution of the protein-antibody binding interface contact areas (100x Å^2^) for Spike-TN1 (purple) and thrombopoietin-TN1 (green) in the molecular dynamics trajectories for PDB templates 1V7M (left) and 1V7N (right). Interaction matrices showing hydrogen bond contribution during the last 50 ns of 200 ns simulations between amino acid residue pairs ordered according to their hydrogen-bond occupancies for the (**b, d**) hTPO-TN1 and (**c, e**) Spike-TN1 complexes for PDB template 1V7M and 1V7N, respectively. Residues belonging to TQLPP are colored in purple and positions for hTPO are based on the PDB template. TN1 Fab residues from heavy and light chains are shaded blue and yellow, respectively.

**Figure S5.**
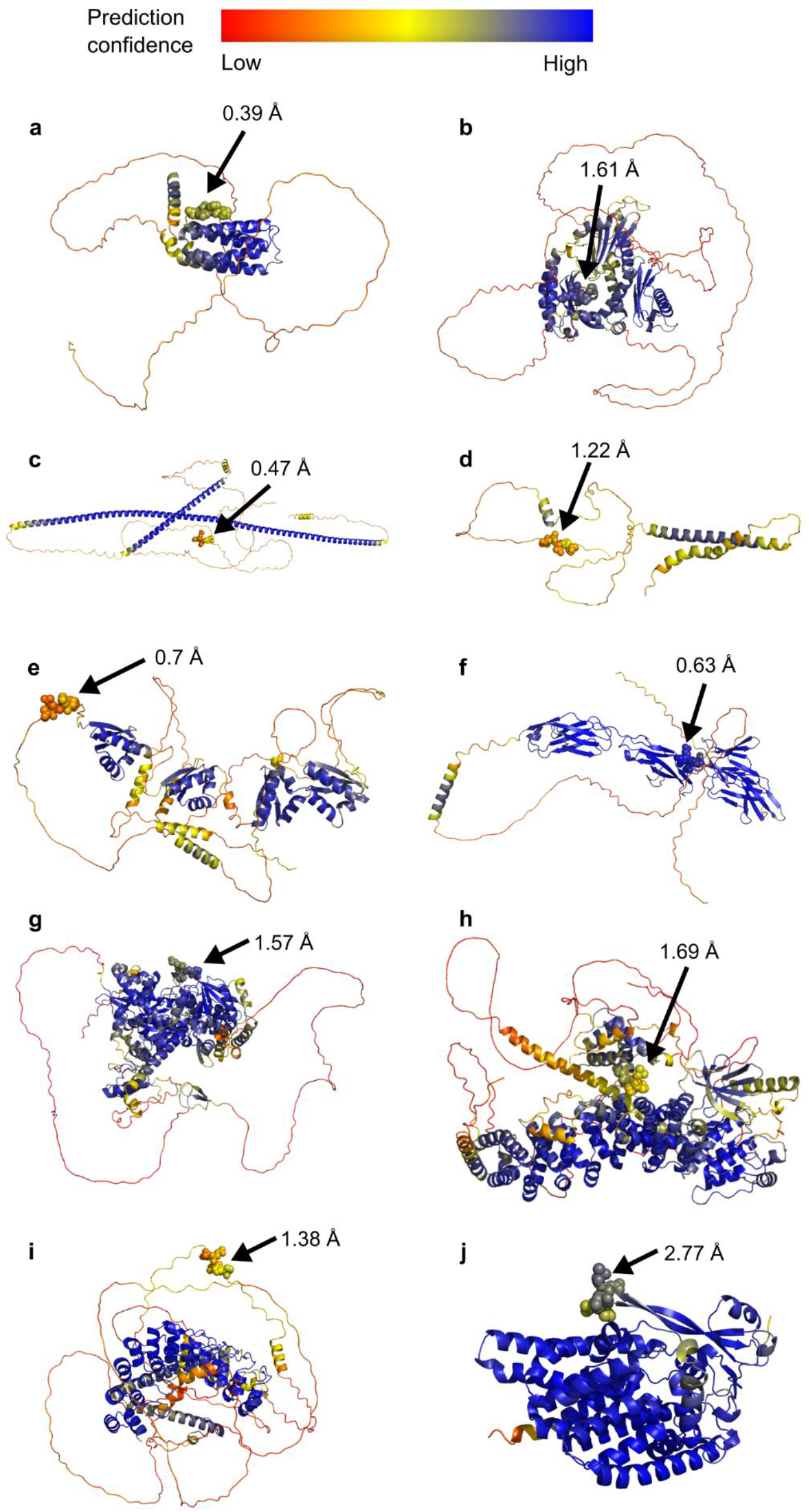
TQLPP motif for 10 human proteins modeled by AlphaFold2. Protein structure models are colored by AlphaFold confidence estimate according to the color bar where red = 25 (low confidence) and blue = 100 (high confidence). TQLPP motif is shown as spheres. RMSD for human TQLPP in the 10 proteins compared to SARS-CoV-2 Spike (PDB id: 6XR8, chain A) is shown. The proteins are (**a**) thrombopoietin (Uniprot: P40225), (**b**) Hermansky-Pudlak syndrome 4 protein (Uniprot: Q9NQG7), (**c**) coiled-coil domain containing protein 85 (Uniprot: Q8N715), (**d**) transmembrane protein 52 precursor (Uniprot: Q8NDY8), (**e**) far upstream element-binding protein 1 (Uniprot: Q96AE4), (**f**) Fc receptor-like protein 4 (Uniprot: Q96PJ5), (**g**) DNA annealing helicase and endonuclease ZRANB3 (Uniprot: Q5FWF4), (**h**) serine/threonine-protein kinase NEK10 (Uniprot: Q6ZWH5), (**i**) espin (Uniprot: B1AK53), and (**j**) ALG12 (Mannosyltransferase ALG12 homolog, Uniprot: Q9BV10). Structural visualization generated with PyMOL (Schrödinger, 2015).

**Figure S6.**
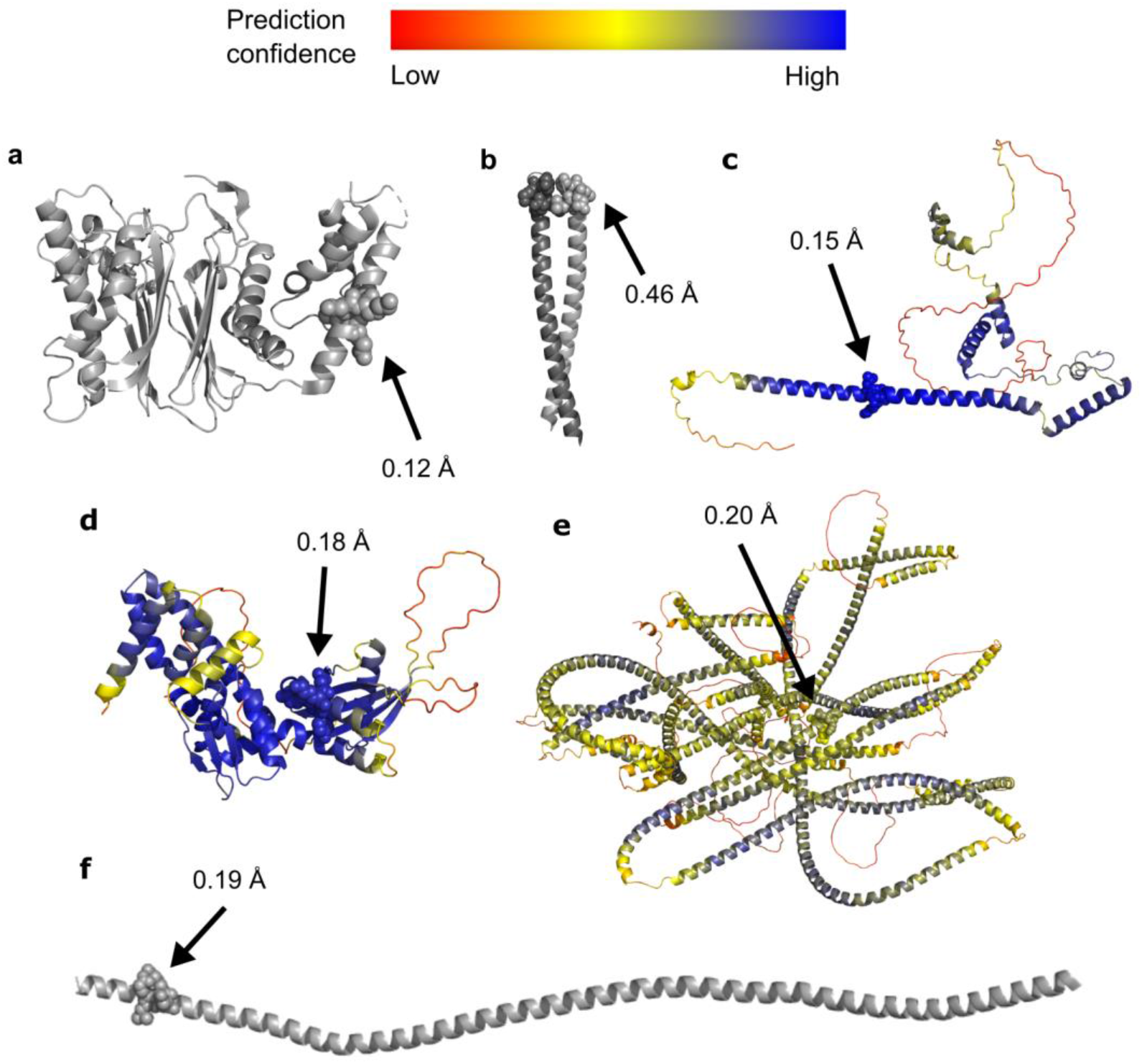
Structure of ELDKY motif for 5 human proteins. Protein structures from PDB are colored gray while AlphaFold2 3D models are colored by AlphaFold confidence estimate according to the color bar where red = 25 (low confidence) and blue = 100 (high confidence). ELDKY motif is shown as spheres. RMSD for human ELDKY in the 5 proteins compared to SARS-CoV-2 Spike (PDB id: 6XR8, chain A) is shown. The proteins are (**a**) protein phosphatase 1A (PDB id: 3FXJ), (**b**) leucine zipper domain of cGMP-dependent protein kinase 1 (PDB id: 3NMD), (**c**) protein FAM228B (Uniprot: P0C875), (**d**) protein Njmu-R1 (Uniprot: Q9HAS0), (**e**) thyroid receptor interacting protein 11 (Uniprot: Q15643), and (**f**) tropomyosin alpha-1 (PDB id: 6X5Z). Structural visualization generated with PyMOL (Schrödinger, 2015).

**Table S1.** RefSeq Select human isoforms that contain pentapeptides found in the 3D-mimics and AF-3D-mimics for SARS-CoV-2 Spike.

**Table S2.** RMSD values resulting from the alignment of the TQLPP region of 1V7M chain X and 1V7N chain X against the TQLPP region of 60 Spike structures. Sorted by RMSD.

| 1V7M X |  | 1V7N X |  |
| --- | --- | --- | --- |
| Spike Structure | RMSD | Spike Structure | RMSD |
| 6ZGG A | 0.36 | 7DCC E | 0.21 |
| 6ZGG B | 0.40 | 7DCC I | 0.27 |
| 7BNN B | 0.43 | 7DCC K | 0.28 |
| 6ZGG C | 0.44 | 7BNN B | 0.42 |
| 7DCC E | 0.44 | 7BNM C | 0.44 |
| 7DCC I | 0.48 | 7BNM B | 0.44 |
| 7DCC K | 0.48 | 7BNM A | 0.44 |
| 7BNM A | 0.52 | 7A25 C | 0.46 |
| 7BNM B | 0.52 | 6XR8 A | 0.46 |
| 7BNM C | 0.52 | 6ZGE C | 0.47 |
| 7A25 C | 0.59 | 6ZGE A | 0.47 |
| 6ZGE A | 0.60 | 6ZGE B | 0.48 |
| 6ZGE B | 0.60 | 6ZGG A | 0.49 |
| 6ZGE C | 0.60 | 7KMK B | 0.49 |
| 7BNN A | 0.60 | 7LRT B | 0.49 |
| 6XR8 A | 0.61 | 6ZGG C | 0.49 |
| 7A25 A | 0.63 | 7A25 A | 0.50 |
| 7LRT B | 0.66 | 6XR8 B | 0.51 |
| 7LRT C | 0.66 | 7LRT C | 0.51 |
| 6XR8 B | 0.68 | 6ZGG B | 0.53 |
| 6XR8 C | 0.71 | 6XR8 C | 0.55 |
| 7A25 B | 0.71 | 7A25 B | 0.57 |
| 6ZP2 A | 0.72 | 7LRT A | 0.58 |
| 6ZP2 B | 0.72 | 7N1U A | 0.59 |
| 6ZP2 C | 0.72 | 7KRQ A | 0.61 |
| 7KMK B | 0.72 | 7BNN A | 0.61 |
| 7LRT A | 0.73 | 7KRQ B | 0.62 |
| 7KRQ A | 0.76 | 7E8C A | 0.64 |
| 7KRQ B | 0.76 | 7BNN C | 0.64 |
| 7N1U A | 0.76 | 7KRQ C | 0.66 |
| 7BNN C | 0.78 | 7KMK C | 0.68 |
| 7KRQ C | 0.79 | 7E8C C | 0.71 |
| 7E8C A | 0.82 | 7E8C B | 0.72 |
| 7LQV A | 0.85 | 7N1U C | 0.73 |
| 7KMK C | 0.87 | 7JJI A | 0.75 |
| 7N1U C | 0.87 | 7JJI B | 0.75 |
| 7E8C C | 0.88 | 7JJI C | 0.75 |
| 7LQV C | 0.88 | 7N1U B | 0.76 |
| 7E8C B | 0.90 | 7KMK A | 0.78 |
| 7N1U B | 0.90 | 7LQV A | 0.78 |
| 7LQV B | 0.91 | 7LQV C | 0.80 |
| 7JJI A | 0.92 | 7LQV B | 0.82 |
| 7JJI B | 0.92 | 6ZP2 A | 0.84 |
| 7JJI C | 0.92 | 6ZP2 C | 0.84 |
| 7CWL B | 0.95 | 6ZP2 B | 0.84 |
| 7CWS R | 0.95 | 7MJG B | 0.85 |
| 7KMK A | 0.96 | 7MJG C | 0.93 |
| 7MJG B | 1.02 | 7MJG A | 0.93 |
| 7MJG A | 1.09 | 7CWL B | 0.94 |
| 7MJG C | 1.10 | 7CWS R | 0.94 |
| 7CWL A | 1.17 | 7CWS O | 1.03 |
| 7CWS O | 1.17 | 7CWL A | 1.03 |
| 7C2L A | 1.21 | 7C2L B | 1.26 |
| 7C2L B | 1.21 | 7C2L C | 1.26 |
| 7C2L C | 1.21 | 7C2L A | 1.26 |
| 7N1Q B | 1.62 | 7N1Q B | 1.56 |
| 7CWL C | 1.67 | 7CWS Q | 1.57 |
| 7CWS Q | 1.67 | 7CWL C | 1.57 |
| 7N1Q A | 1.68 | 7N1Q A | 1.61 |
| 7N1Q C | 1.71 | 7N1Q C | 1.64 |

**Table S3.** RMSD values resulting from the alignment of the TQLPP region from 60 Spike structures and three modeled states, representing a conformational ensemble of TQLPP in Spike, sorted by RMSD. **Separate Excel sheet**

**Table S4.** Distribution of MaSIF values. **Separate Excel sheet**

**Table S5.** Statistical comparison of MaSIF binding scores for antibody complexes.

| **MaSIF binding scores** |  |  |  |
| --- | --- | --- | --- |
| **Comparison** |  | **p-value** | **Significant<sup>1</sup>** |
| **Random** | Spike-Ab | 4.83E-75 | Yes |
| **Random** | hTPO-TN1 | 3.00E-08 | Yes |
| **Random** | Spike-TN1 | 5.24E-26 | Yes |
| **Spike-Ab** | hTPO-TN1 | 3.37E-02 | No |
| **Spike-Ab** | Spike-TN1 | 7.68E-09 | Yes |
| **hTPO-TN1** | Spike-TN1 | 1.92E-04 | Yes |

<sup>1</sup> Compared to Bonferroni corrected p-value (<8.33E-03) for alpha = 0.05

**Table S6.**
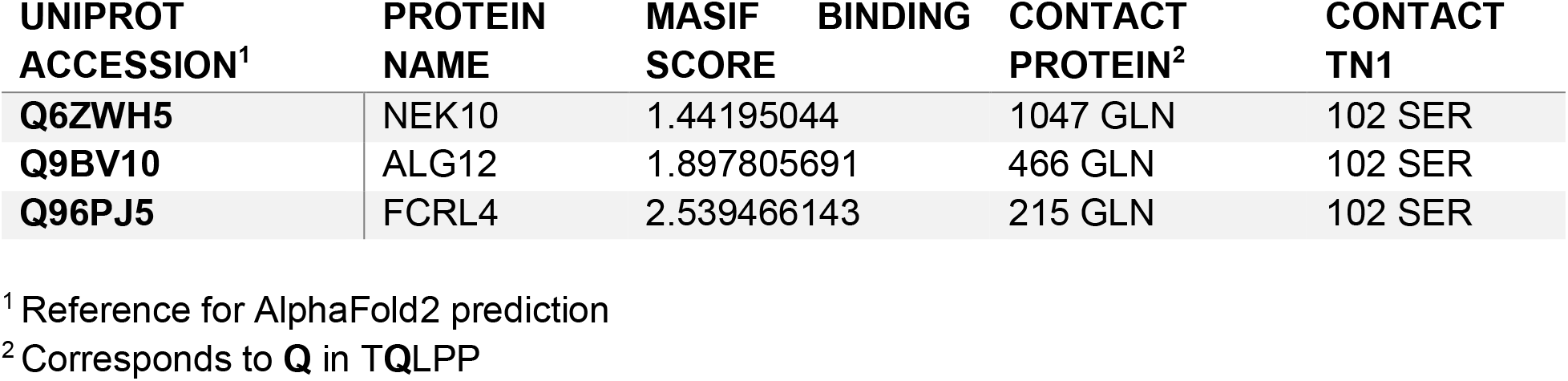
MASIF binding scores of other human proteins in complex with TN1.

